# SOX21 Suppresses Glioblastoma Growth by Repressing AP-1 Activity

**DOI:** 10.1101/2024.07.05.601643

**Authors:** Eltjona Rrapaj, Juan Yuan, Idha Kurtsdotter, Vsevolod Misyurin, Guido Baselli, Oscar Persson, Maria Bergsland, Jonas Muhr

**Author notes:** Corresponding author: Jonas Muhr. ^#^ These authors contributed equally to this work (Eltjona Rrapaj, Juan Yuan). The authors declare no potential conflicts of interest.

## Abstract

**Background:** Treatment-resistant glioblastoma stem cells (GSCs) drive glioblastoma (GBM) growth and recurrence. Thus, targeting the molecular machinery that sustains GSCs in an undifferentiated and self-renewing state is a promising therapeutic strategy. The transcription factor SOX21 effectively suppresses the tumorigenic capacity of GSCs. However, the mechanism by which SOX21 impedes GSC features is unknown.

**Methods:** Patient-derived GSCs were engineered with a transgenic TetOn system to enable inducible expression of SOX21 or appropriate controls. The capacity of SOX21 to incapacitate GSCs was assessed using *in vitro* cell culture models and orthotopic mouse models. Cellular and genome-wide techniques, including RNA-seq, ChIP-seq, and ATAC-seq, were employed to examine the mechanisms by which SOX21 regulates GSCs.

**Results:** We show that SOX21 expression in primary GSCs induces an anti-tumorigenic transcriptional program, aligning with clinical data showing a positive correlation between SOX21 levels and improved GBM patient survival. Induced SOX21 expression in GSCs within pre-established GBM reduces their capacity to sustain tumor growth and significantly extends the survival of the transplanted mice. Mechanistically, SOX21 functions as a tumor suppressor by binding a large set of AP-1-targeted chromatin regions, leading to epigenetic repression of AP-1-activated genes that support GSC survival and proliferation. Consistently, the anti-tumorigenic activities of SOX21 are replicated by AP-1 inhibitors, while overexpression of the AP-1 family member, c-JUN, counteracts these effects.

**Conclusion:** Our findings identify SOX21 as a key regulator that prevents GSC malignancy by targeting and repressing an AP-1-driven, tumor-promoting gene expression program. These results highlight SOX21-regulated pathways as promising therapeutic targets for GBM.

**Key Points:**

- Induced SOX21 expression suppresses GSCs and inhibits the growth of established GBM
- SOX21 acts as a tumor suppressor in GSCs by directly repressing AP-1-driven genes
- Pharmacological inhibition of AP-1 mimics SOX21 activity in GSCs

**Importance of the Study:** GBM is the most common and aggressive malignant brain tumor in adults. Recurrence following treatment often stems from the failure of therapeutic interventions to effectively target GSCs, which serve as the primary reservoir for tumor regrowth. The resilience of GSCs to treatment is partly due to the inactivation of intrinsic tumor suppressor programs that would otherwise direct GSCs to cellular senescence and death. This study demonstrates that increased expression levels of the tumor suppressor SOX21 in pre-established GBM disrupt tumor progression by disabling self-renewing GSCs. We show that SOX21 exerts its tumor-suppressive function by targeting and repressing an AP-1-driven gene network, which is a key regulator of GSC maintenance and proliferation. By uncovering the molecular mechanisms through which SOX21 controls GSC biology, our findings provide valuable insights for basic cancer research and may inform the development of novel therapeutic strategies for GBM.

## Introduction

GBM is the most prevalent and aggressive form of primary brain tumors^1^. It is characterized by uncontrolled growth, cellular and molecular heterogeneity, and a high propensity for recurrence. One major reason why conventional therapies unavoidably fail is explained by the presence of treatment-resistant GSCs^2^. These cells exhibit extensive self-renewing capacity, supporting rapid tumor growth, and high cellular plasticity, which contributes to inter- and intra-tumoral diversification^2,3^. Additionally, GSCs possess potent tumor-initiating potential^4^, underscoring the need for novel therapeutic strategies that target these cells to curb GBM progression and recurrence.

Stem cells, both in normal and malignant contexts, are regulated by an interplay of transcription factors that either maintain cells in an undifferentiated, self-renewing state or facilitate cellular differentiation and cell-cycle exit. Several transcription factors that promote the growth of healthy stem cells, including members of the *SOX*, *POU*, *AP-1*, *GLI,* and *E2F* families, have also been implicated in GSC maintenance and tumorigenicity^5–10^. For instance, SOX2, a key regulator of stem cells in developing and adult tissues, is also critical for GSC propagation and their tumorigenic capacity^11,12^. Similarly, AP-1 family members, which have ubiquitous roles in regulating cell proliferation and survival^13^, have been shown to promote stem cell-like properties and aggressiveness of GSCs^8,9^.

In contrast, the transcription factor SOX21 exhibits the opposite activity and reduces neural stem cell proliferation and promotes their differentiation^14^. Notably, SOX21 expression is significantly lower in high-grade gliomas compared to low-grade tumors^15^. Consistent with this, forced expression of SOX21 in human GSCs diminishes their ability to form secondary tumors following orthotopic transplantation into mice^15,16^. Conversely, genetic ablation of *Sox21* in mouse brain stem cells strongly increases their propensity to form GBM-like tumors in an H-RAS/AKT-driven glioma model^15^. These gain- and loss-of-function experiments underscore the capacity of SOX21 in preventing oncogenic transformation of brain stem cells and demonstrate that forced expression of SOX21 in GSCs before transplantation curbs their capacity to initiate tumor formation in mice. However, from a therapeutic perspective, it is critical to examine whether SOX21 induction in already established GBM can block GSC propagation and thereby inhibit further tumor growth. Additionally, the molecular mechanisms by which SOX21 counterbalances genetic pathways that preserve GSC properties and promote GBM progression remain largely unexplained.

In this study, we generated an inducible expression system in patient-derived GSCs to investigate the therapeutic potential of SOX21. We demonstrate that induced SOX21 expression, but not SOX2 expression, suppresses GSC propagation, which counteracts further tumor growth of established GBM and significantly improves the survival of the orthotopically transplanted mice. AP-1 proteins promote GSC self-renewal and survival. We now show that SOX21 and AP-1 proteins interact and that a major portion of SOX21 binding is directed to chromatin regions targeted by the AP-1 family member c-JUN, resulting in repression of AP-1 activated genes and as a result an inhibition of GSC proliferation and survival.

## Materials and Methods

See Supplementary Methods for additional details.

### SOX21 Expressing GSCs

Human glioblastoma (GBM) samples and primary GBM cell cultures were obtained from Karolinska Institute Biobank (Ethical permit 2023-01366-01). Cells were cultured on poly-L-Ornithine (Sigma Aldrich) /Laminin (Sigma Aldrich)-coated plates in Human NeuroCult NS-A Proliferation media (STEMCELL Technologies) supplemented with 2 μg/mL heparin (STEMCELL Technologies), 10 ng/mL FGF (STEMCELL Technologies), 20 ng/mL EGF (STEMCELL Technologies), and penicillin/streptomycin (Gibco).

SOX21 and SOX2 expression was induced via DOX (Sigma Aldrich)-regulated lentiviral transduction (pLVX-SOX21, pLVX-SOX2, or pLVX). Selection with 6 μg/mL Puromycin (Gibco) lasted four weeks. For in vivo experiments, cells were further transduced with a constitutive Luciferase-expressing lentivirus (pGK-Luc). All primary cells were passaged below 10 and tested for mycoplasma using MycoAlert Mycoplasma Detection kit (Lonza). DOX (200 ng/mL) was added for 48 hours pre-assay unless otherwise specified.

### Intracranial Transplantation of GSCs

Animal experiments adhered to Swedish animal welfare laws (Dnr 3696-2020). Human GBM cells (150,000 per injection) were prepared at 50,000 cells/μL with >80% viability and maintained on ice for up to two hours before transplantation. Stereotactic injections targeted the striatum in immunocompromised NOD.CB17-PrkcSCID/J mice (6-10 weeks old) at 2.0 mm lateral, 1.0 mm anterior, and 2.5 mm depth from bregma (3 µl). Pre-operative analgesia (0.1 mg Norocarp/Carprofen (Zoetis), 2-5 mg/kg) was administered subcutaneously, and anesthesia was maintained with 2-3% isoflurane (Zoetis) in O2 (1 L/min) during transplantation. Local analgesia (2.5 mg/mL, Marcain, Aspen Pharmacare) was applied at the incision site. Postoperative care included subcutaneous administration of 0.001 mg Temgesic/Vetergesic (0.05-1 mg/kg) (Ceva Santé Animale) for two days, with daily monitoring for one week. Mice exhibiting distress were euthanized according to ethical guidelines. Survival data were recorded for statistical analysis.

### In Vivo Bioluminescence Imaging

Tumor progression was assessed using IVIS Spectrum CT Imaging (Perkin Elmer). Mice received intraperitoneal injections of 150 mg/kg D-Luciferin (Promega) and were imaged 23 minutes post-injection to capture peak luminescence. Animals were maintained at 37°C under gaseous anesthesia (2-3% isoflurane in 1 L/min O2) during imaging. Data were processed in Living Image software (Perkin Elmer).

### Tissue Processing and Histology

Mice were deeply anesthetized with avertin (Sigma Aldrich) before transcardial perfusion with PBS (Gibco) followed by 4% paraformaldehyde (Sigma Aldrich). Brains were extracted, post-fixed in PFA, and processed for cryopreservation or paraffin embedding. Coronal sections (5-15 μm) were prepared. Tumor morphology was assessed via hematoxylin and eosin (H&E) staining (Histolab). Images were acquired using a Zeiss AxioScan.Z1 slide scanner and analyzed with QuPath (v0.2.3) and ImageJ.

### Immunofluorescence

Tissue sections underwent deparaffination, antigen retrieval (Dako) and blocking with 4% donkey serum (Jackson ImmunoResearch)/FBS. Primary antibodies (SOX21, SOX2, HuNu, Ki67) were incubated overnight at room temperature, followed by secondary antibody labeling. Nuclei were counterstained with DAPI. Images were acquired using a ZEISS LSM700 confocal microscope and analyzed with ImageJ.

### Neurosphere Formation Assay

Primary GSCs were cultured in low-attachment plates with DMEM/F12-Glutamax (Gibco), N-2 (Gibco), B-27 (Gibco), recombinant human EGF, bFGF, 1.5% FBS (Cytiva), and 0.05% BSA (Sigma Aldrich). Rock-inhibitor/Y-27632 (1 µM, STEMCELL Technologies) was added for the first 1-2 days. DOX (250 ng/mL) or AP-1 inhibitors (10 μM SR11302 (ChemCruz), 100 μM T5224 (Cyman Chemical)) were applied on day three. Medium was refreshed every two days. Spheroid growth was assessed after 10 days (DOX) or 14 days (AP-1 inhibitors) using a Leica CTR 4000 microscope and ImageJ.

### Cell Imaging and Assays

Cells were imaged using the EVOS cell imaging system (10x magnification). Proliferation was measured with a Click-iT EdU Flow Cytometry Assay Kit (Invitrogen) and analyzed by flow cytometry (BD Canto II, 50,000 events/sample). Apoptosis was assessed using FITC Annexin V detection (BD Biosciences) and analyzed by FACS (BD FACSCanto II). Data were processed using FlowJo software.

### Western Blot and Co-IP

Protein lysates were prepared using RIPA buffer (Sigma Aldrich) supplemented with a protease inhibitor mixture (Roche). Cells were lysed on ice for 30 min with intermittent vortexing, followed by centrifugation at 14,000 × g for 10 min at 4 °C to clear debris. Protein concentrations were determined using a Bradford Assay (Bio-Rad).

For Western blot, equal amounts of protein lysates were mixed with loading buffer, denatured at 95°C for 5 min, and resolved on an 8–12% SDS-PAGE gradient gel alongside a protein ladder (ThermoScientific). Proteins were transferred onto a nitrocellulose membrane (Amersham) and blocked in 5% milk in PBS/0.1% Tween-20 (Sigma) for 1 h. Membranes were incubated overnight at 4°C with primary antibodies (SOX21, SOX2, FLAG, HA, Ki67, H3K27Ac, H3K4me1) at optimized dilutions. Secondary HRP-conjugated antibodies (Vector Laboratories) were applied at a 1:5,000 dilution, followed by detection with ECL substrate (ThermoScientific).

For Co-IP, cells were lysed in a buffer containing 50 mM Tris-HCl (pH 7.4), 150 mM NaCl, 1% NP-40, and protease/phosphatase inhibitors. Lysates were cleared by centrifugation at 12,000 × g for 15 min at 4°C. 50 U of DNase I (ThermoScientific) per 1 mL of lysate was added and incubated at 37°C for 20 min. Immunoprecipitation was performed with 5 µg anti-HA or control IgG conjugated to Dynabeads (ThermoScientific) overnight at 4°C. Beads were washed three times, and bound proteins were eluted with SDS sample buffer. Proteins were separated by SDS-PAGE and detected by immunoblotting using anti-FLAG and anti-HA antibodies to assess interactions between SOX21 and c-JUN.

Primary antibodies used were SOX2 (SEVEN HILLS BIOREAGENTS), SOX21 (R&D), FLAG (Abcam; Sigma Aldrich), HA (Santa Cruz), mouse IgG (Millipore).

Secondary antibodies used were Donkey anti Rabbit HRP, Donkey anti Mouse HRP, Donkey anti Goat HRP (Jackson ImmunoResearch).

### Statistical Analysis

GraphPad Prism (v9.0) was used for all statistical analyses. Significance was determined using unpaired t-tests or one-way ANOVA. Kaplan-Meier survival curves were analyzed using log-rank (Mantel-Cox) tests. For gene set enrichment, hypergeometric distribution (phyper, R software) was used. A p-value < 0.05 was considered significant. For Sox-21 expression analysis in TCGA glioblastoma cohort, RNA sequencing data and overall survival of 151 patients with Glioblastoma, were downloaded from The Cancer Genome Atlas (TCGA) data portal51 (TCGA; https://cancergenome.nih.gov/). The cut-off value for SOX21 mRNA expression was determined based on its median value. The relationship between SOX21 mRNA expression and overall survival was analyzed using the log-rank test with R software (version 4.3.1; R Foundation for Statistical Computing, Vienna, Austria)^17^ .

### Data Availability

Raw sequencing data are accessible via BioProject ID PRJNA1128363.

## Results

### SOX21 Expression is Confined to GSCs and Correlates With Improved Patient Survival

To further explore the role of SOX21 in GBM, we examined its protein expression pattern in surgical GBM specimens (IDH wild type) using immunofluorescence. In these tumors, SOX21 expression was largely restricted to SOX2^+^ cells (Figure 1A-C, G; Supplementary Figure S1A-D). Additionally, SOX21 expression was detected in most Ki67^+^ proliferating cells (Figure 1D-G; Supplementary Figure S1A-D). Hence, in GBM the expression of SOX21 is enriched in the GSC compartment.

**Figure 1.**
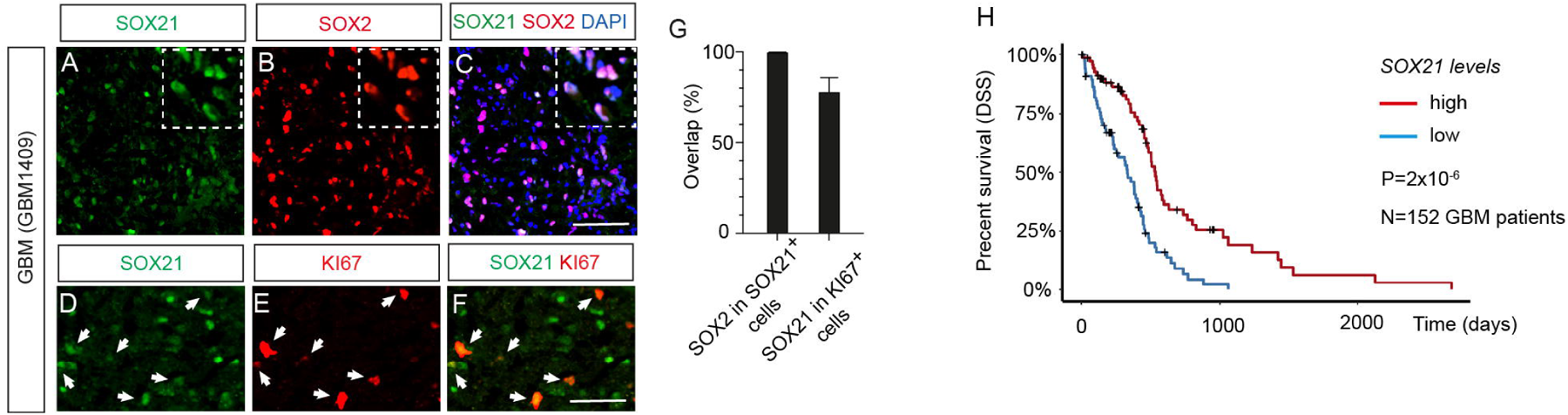
*SOX21* Expression Levels Hold Prognostic Value for GBM Patient Survival. (A-C) Immunofluorescence analysis of a GBM surgical sample (GBM1409) reveals SOX21 protein localization (green in A and C), predominantly in *SOX21* cells (red in B and C). Insets show magnified views of the tumor area. Scale bar, 50 µm. (D-F) SOX21 expression (green in D and F) can be detected in most actively proliferating Ki67^+^ cells (red in E and F). Scale bar, 50 µm. (G) Statistical co-expression analysis of SOX21, SOX2, and Ki67 proteins in the GBM1409 sample. (H) Kaplan-Meier survival curves derived from TCGA GBM mRNA expression data (n = 152 samples) demonstrate significantly improved disease-free survival in patients with high SOX21 expression (red line) compared to those with low expression (blue line) (p = 2 × 10⁻⁶).

Previous analyses of gene expression data sets from low-grade (WHO grade II and III) and high-grade (WHO grade IV, GBM) glioma have revealed an inverse correlation between SOX21 expression levels and glioma malignancy^15^. To examine if the expression levels of *SOX21* also hold a prognostic value for GBM patient survival, we divided the GBM mRNA data sets from “The Cancer Genome Atlas” (TCGA) into two groups, based on *SOX21* levels, and correlated these to the patient survival. Notably, Kaplan-Meier survival analyses revealed that patients with high *SOX21* expression exhibited significantly better Disease-Specific Survival (DSS) (p= 2x10^-6^), compared to those with low *SOX21* expression (Figure 1H).

### Elevated SOX21 Levels Suppress GSC Proliferation and Survival

The positive correlation between *SOX21* expression levels and patient survival suggests that high levels of SOX21 may exert an anti-tumorigenic function in high-grade gliomas. To address this possibility, we generated a tetracyclin-inducible (TetOn) lentiviral vector encoding a FLAG-tagged version of *SOX21* (*pLVX-SOX21*) and, as a control, a version lacking insert (*pLVX*) (Figure 2A). The resulting viruses were stably transduced into primary GSCs derived from three different GBM tumors (Supplementary Table S1); GSCs hereafter referred to as JM11, JM12, and JM13. In comparison to 209 glioma samples of known molecular subtypes^18,19^, RNA-seq analysis revealed that these GSCs were best classified as classical (JM11), proneural (JM12), and mesenchymal (JM13) subtypes, respectively (Supplementary Figure S2). Following selection for successfully transduced cells, the expanded GSCs were confirmed, with immunoblotting and immunofluorescence, to specifically upregulate SOX21 upon doxycycline (DOX) treatment (Figure 2B; Supplementary Figure S3A, B).

**Figure 2.**
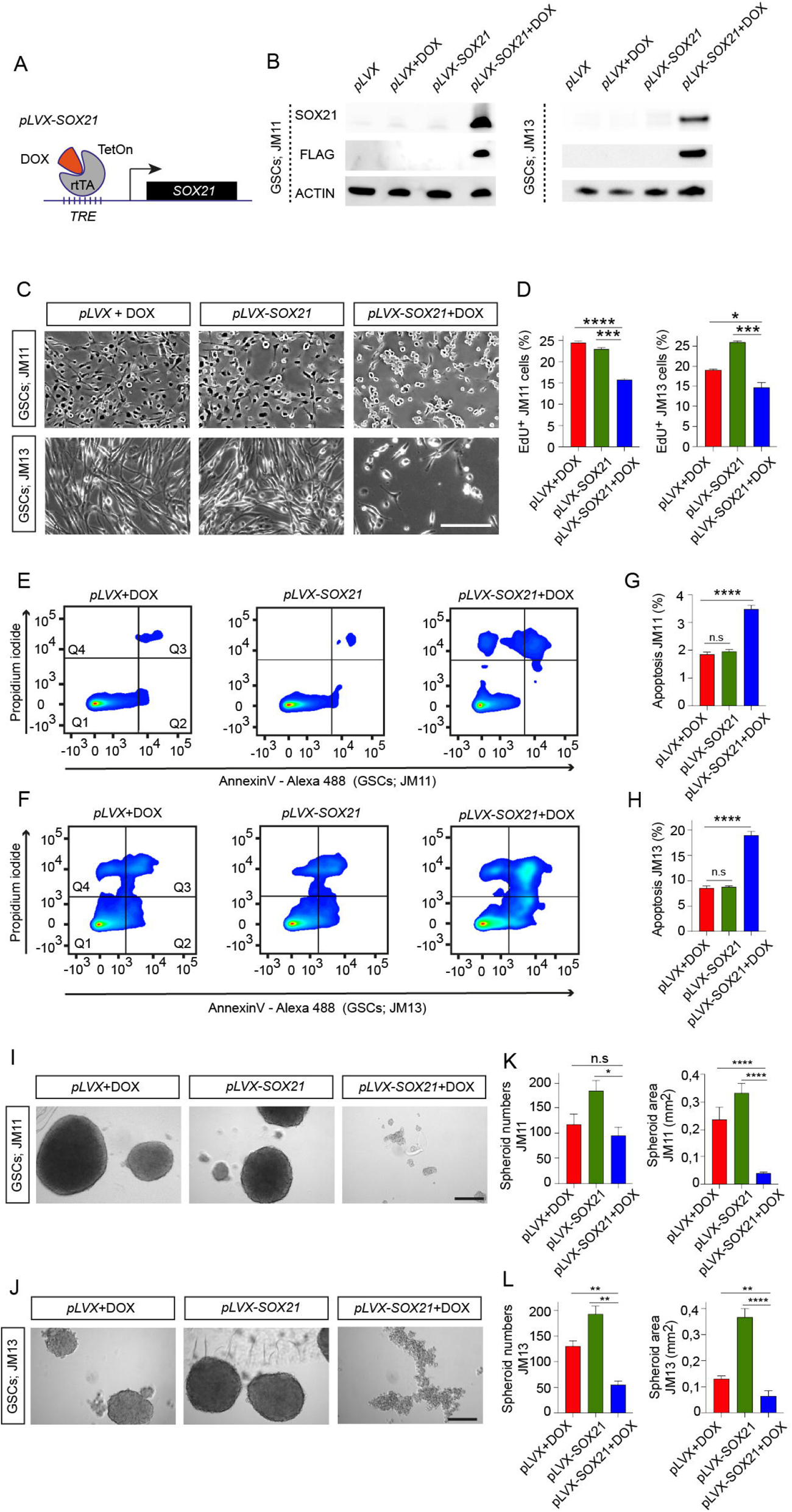
Effects of Inducible SOX21 Expression on GSC Growth. (A) Schematic representation of the tetracycline-inducible (Tet-On) SOX21 expression system. rtTA, tetracycline-controlled transactivator; TRE, tetracycline response element; DOX, doxycycline. (B) Western blot analysis of FLAG-tagged SOX21 expression in GSC lines (JM11 and JM13) transduced with either control (pLVX) or inducible SOX21 expression vectors (pLVX-SOX21), cultured with or without DOX for 48 hours. (C) Representative images showing reduced cell density in GSC monolayer cultures following DOX-induced SOX21 expression. Scale bar, 70 µm. (D) Quantification of EdU incorporation in control and SOX21-expressing GSCs (JM11 and JM13). (E, F) Flow cytometry analysis of JM11 (E) and JM13 (F) GSCs transduced with control (pLVX) or SOX21-inducible (pLVX-SOX21) vectors and cultured with or without DOX for 4 days. Quadrant distributions: Q1 (live cells), Q2 (early apoptotic cells), Q3 (late apoptotic cells), and Q4 (necrotic cells). (G, H) Quantification of flow cytometry experiments showing increased apoptosis in GSCs following DOX-induced SOX21 expression, as assayed by Annexin V labeling. (I, J) Representative images illustrating the sphere forming capacity of JM11 (I) and JM13 (J) GSCs with or without SOX21 induction. Scale bar, 500 µm. (K, L) Bar graphs quantifying the number and size of spheroids formed by JM11 (K) and JM13 (L) GSCs.

DOX-induced SOX21 expression, “*pLVX-SOX21*+DOX”, significantly reduced GSC density in monolayer cultures compared to controls, “*pLVX*+DOX” and “*pLVX-SOX21”* (non-Dox) (Figure 2C). Additionally, the incorporation of the thymidine analog EdU revealed a marked decrease in the proportion of GSCs undergoing active proliferation upon SOX21 induction (Figure 2D). Flow cytometry (FACS) based analysis further demonstrated that SOX21 upregulation significantly increased the proportion of early and late apoptotic GSCs, as evidenced by Annexin V expression and propidium iodide labeling (Figure 2E-H). To assess the impact of SOX21 on GSC clonal expansion capabilities, we performed sphere-forming assays. Within 5-7 days, DOX-induced SOX21 expression resulted in a notable reduction in both the number and size of the tumor spheres formed, compared to control conditions (Figure 2I-L). Together, these findings indicate that increased levels of SOX21 suppress GSC proliferation, enhance apoptosis, and limit clonal expansion potential, supporting its potential as a tumor suppressor in GSCs.

### Induced SOX21 Expression in Established GBM Suppresses Tumor Progression

To further evaluate the anti-tumorigenic effects of SOX21, we investigated its capability to restrict GBM growth *in vivo*. Previous studies have demonstrated that viral-mediated SOX21 expression in GSCs and glioma cell lines inhibits their tumor-initiating capacity in orthotopic mouse models^15,16^. However, from a therapeutic perspective, it is critical to determine whether SOX21 induction in established GBM can block GSCs to promote further tumor growth. To address this issue, we transplanted GSCs of distinct molecular subtypes (JM11, JM12, and JM13) – engineered with either DOX-inducible SOX21 expression constructs or control vectors, along with a constitutive lentiviral Luciferase (Luc) reporter – into the striatum of NOD-SCID mice (Figure 3A). Serving as an additional control, we also transplanted mice with Luc-expressing GSCs additionally carrying a DOX-inducible SOX2 expression system (*pLVX-SOX2*; Supplementary Figure S3C). Following tumor establishment, confirmed via luciferase activity measurements using an *in vivo* imaging system (IVIS), mice were provided with DOX-supplemented food to induce transgenic expression^20^ (Figure 3A, B; Supplementary Figure S4A).

**Figure 3.**
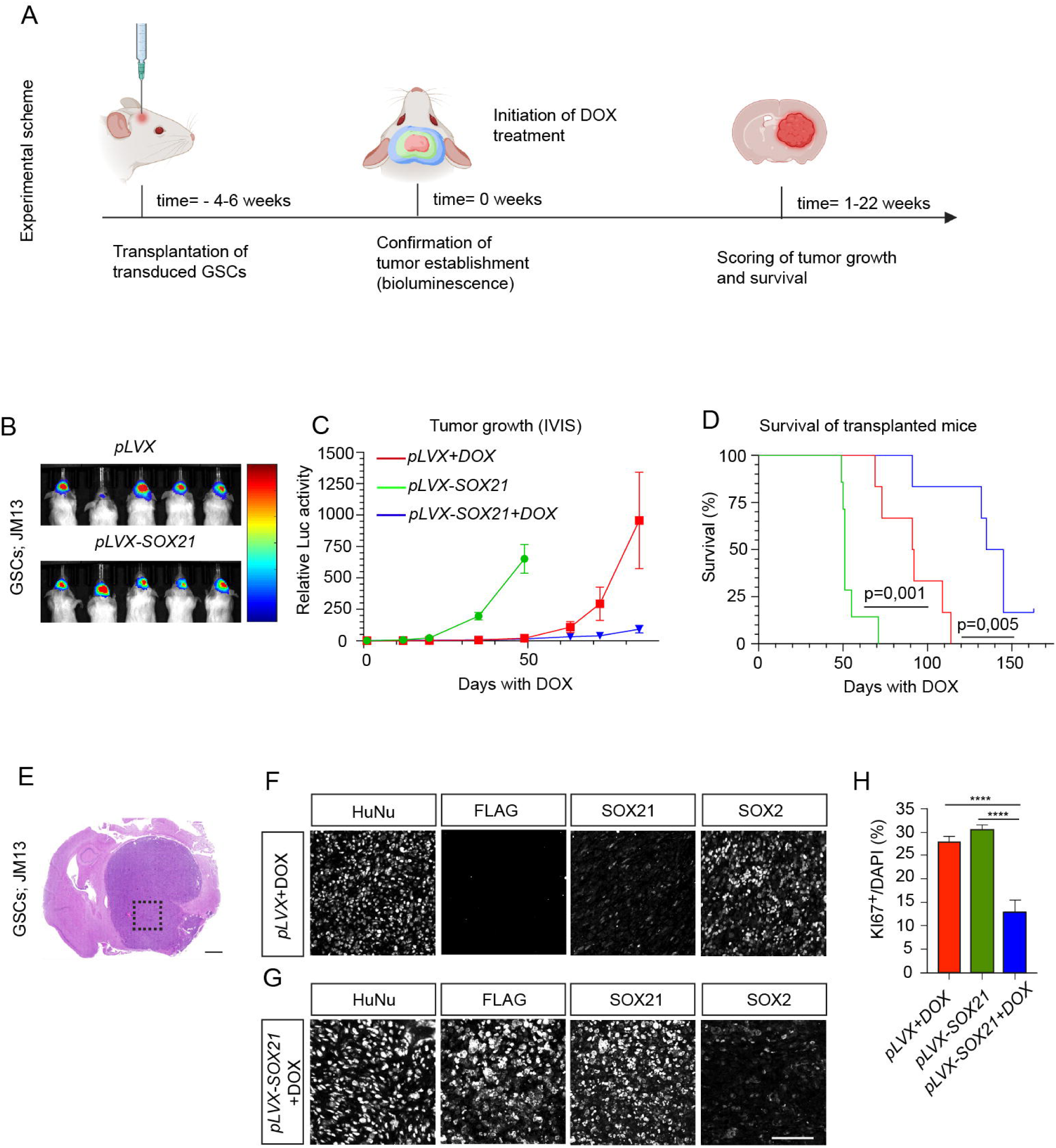
Induced SOX21 Expression Suppresses Growth of Pre-Established GBM in Mice. (A) Experimental timeline illustrating the assessment of DOX-induced SOX21 expression in pre-established GBM tumors, monitored via bioluminescence imaging. (B) Bioluminescence images confirm tumor establishment before the initiation of DOX treatment in mice transplanted with control or SOX21-inducible GSCs (JM13). (C) Tumor growth curves following control or DOX supplementation. The tumor burden quantified by luciferase intensity. (D) Kaplan-Meier survival analysis demonstrates that SOX21 induction significantly prolongs survival in tumor-bearing mice. (E) Representative H&E-stained section of an end-stage tumor from a DOX-treated mouse transplanted with control GSCs (JM13). Scale bar, 1,5 mm. (F, G) High-magnification immunohistochemical images showing expression of human nuclei (HuNu), FLAG-tagged SOX21, endogenous SOX21, and SOX2. Scale bar, 75 µm. (H) Quantification of Ki67-expressing cells demonstrates a reduction in proliferative cells upon SOX21 expression.

Bioluminescence imaging every two weeks revealed that SOX21 induction (*pLVX-SOX21*+DOX) significantly reduced GBM progression compared to control tumors (*pLVX*+DOX and *pLVX-SOX21*) (Figure 3C; Supplementary Figure S4B). The suppression of tumor growth was further reflected by the significantly prolonged mean survival time for SOX21-induced mice relative to controls (Figure 3D; Supplementary Figure S4C, D). Immunohistochemical analysis of end-stage tumors confirmed that most cells retained transgene expression (Figure 3E-G; Supplementary Figure S4E-G). Moreover, exogenous SOX21 expression in GBM significantly decreased the proportion of Ki67^+^ proliferative cells and SOX2^+^ GSCs (Figure 3F-H; Supplementary Figure S4F-H). These findings demonstrate that SOX21 expression in pre-established GBM suppresses tumor progression by depleting proliferative GSCs. In contrast, DOX-induced expression of SOX2 had no significant effect on tumor growth or transplanted mice’s mean survival outcome (Supplementary Figure S4I-N).

### SOX21 Represses Tumor-Promoting Genes and Shares Chromatin Targets with AP-1

To elucidate the molecular mechanisms underlying SOX21-mediated tumor suppression, we profiled its genome-wide binding and transcriptional regulatory effects using RNA-seq and ChIP-seq in two GSC lines (JM11 and JM13), with SOX2 serving as a control. SOX21 induction for 48 hours resulted in the deregulation of over a thousand genes (Figure 4A, B; Supplementary Figure S5A, B; Supplementary Table S2 and S3). Among the most significantly upregulated genes was *CDKN1A*, which encodes the tumor suppressor p21 (Figure 4A, B; Supplementary Figure S5A-C). Gene ontology (GO) analysis revealed that SOX21 enhances the expression of apoptosis-related pathways, including “*neuron apoptotic process”*, *glial cell apoptotic process* ”, but also pathways such as “*negative regulation of cell cycle process* ” (Figure 4C; Supplementary Figure S5D). Conversely, SOX21 downregulated genes involved in glioma progression, such as *CDK6*, *EFNB2*, *HDAC9,* and *SOX2* (Figure 4A, B; Supplementary Figure S5A, B)^21–25^, and genes associated with “*Positive regulation of cell cycle process”*, which further supports its role as a tumor suppressor (Figure 4C; Supplementary Figure S5D). SOX2 overexpression deregulated genes distinct from those altered by SOX21 (Supplementary Figure 5E-H), although it did downregulate the cell cycle gene *CCND1*^26^ (Supplementary Figure S5E, F). Together, these findings highlight SOX21 as a transcriptional regulator that opposes tumor-promoting gene expression in GSCs.

**Figure 4.**
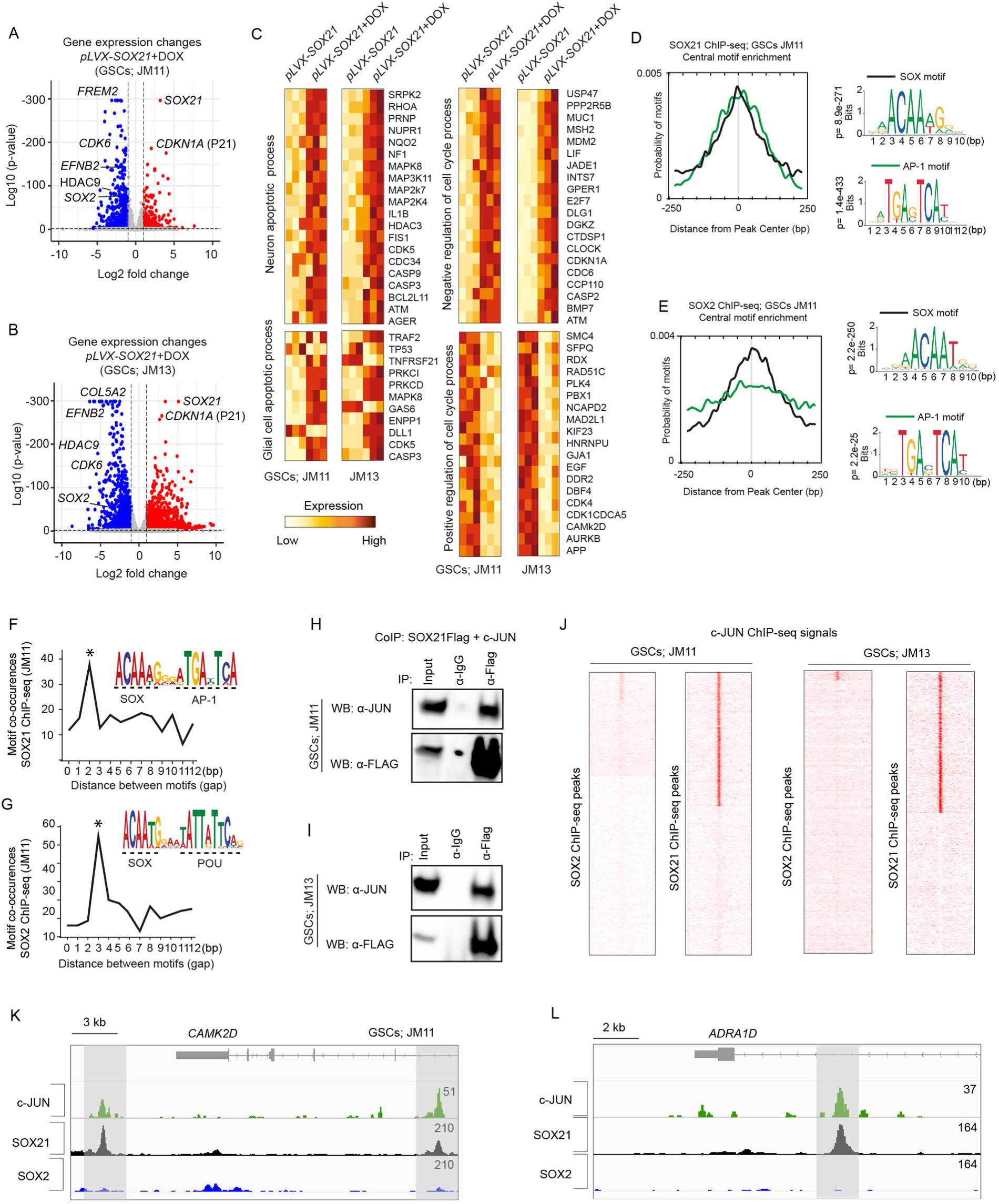
Transcriptomic Impact of SOX21 Expression in GSCs. (A, B) Volcano plots displaying differentially expressed genes in JM11 (A) and JM13 (B) GSCs after 48 hours of DOX-induced SOX21 expression. Genes significantly upregulated (red) and downregulated (blue) are shown (false discovery rate < 0.01). (C) Heatmaps depicting gene sets and associated Gene Ontology (GO) terms deregulated upon SOX21 induction. (D, E) Enrichment analysis of SOX motifs (black line) and AP-1 motifs (green line) within SOX21 (D) and SOX2 (E) ChIP-seq peaks, with motif distances to the center of SOX21 (D) and SOX2 (E) peaks measured in base pairs (bp). P-values of best-matching SOX and AP-1 motifs are shown. (F, G) Graphs illustrating the most significant spacing distances between centrally enriched SOX and AP-1 motifs in SOX21 ChIP-seq peaks (F) and between SOX and POU motifs in SOX2 ChIP-seq peaks (G). The gap between motifs is based on the spacing between the last nucleotide in the SOX motif and the first nucleotide in AP-1 (F) or POU (G) motifs (H, I) Co-immunoprecipitation assays showing interactions between FLAG-tagged SOX21 and c-JUN in GSCs (JM11 and JM13) treated with DNase I. (J) Genomic mapping of c-JUN ChIP-seq reads in JM11 and JM13 within SOX21 ChIP-seq peak regions. (K, L) Genomic regions surrounding CAMK2D (K) and ADRA1D (L) genes, illustrating ChIP-seq binding profiles for c-JUN, SOX21, and SOX2 in JM11 cells.

ChIP-seq experiments revealed thousands of SOX21 and SOX2 binding sites (peaks), with high reproducibility across replicates (Supplementary Figure S6A; Supplementary Table S4). Notably, approximately one-third of SOX21 binding sites were consistent across both JM11 and JM13 GSC lines, whereas SOX21 and SOX2 shared fewer than 10% of their binding sites (Supplementary Figure S6B-D). Interestingly, while centrally located SOX binding motifs were present in both SOX21 and SOX2 peaks (Supplementary Figure S6E, F), motif analysis revealed a significant enrichment of AP-1 transcription factor binding sites in SOX21-bound regions (Figure 4D; Supplementary Figure S6G). This AP-1 motif enrichment was not attributable to cross-reactivity of the antibodies used in the ChIP-seq experiments (Supplementary Figure S6I, J). In contrast, no AP-1 binding motif enrichment was detected at SOX2-bound chromatin regions (Figure 4E; Supplementary Figure S6H).

Furthermore, analysis of SOX21 peak regions revealed a strong co-enrichment of SOX and AP-1 motifs within 1-2 base pairs (Figure 4F; Supplementary Figure S6K). By comparison, SOX2-targeted chromatin regions were preferentially enriched for SOX and POU motifs, spaced 2-3 base pairs apart (Figure 4G; Supplementary Figure S6L). Consistent with this observation, co-immunoprecipitation experiments confirmed a physical interaction between SOX21 and the AP-1 family member c-JUN, a highly expressed AP-1 family member in GSCs (Figure 4H, I; Supplementary Tables S1, 2).To further examine the functional interplay between SOX21 and AP-1 we performed c-JUN ChIP-seq experiments in JM11 and JM13 GSCs. These analyses revealed that approximately 50 % of the SOX21-bound chromatin regions overlapped with c-JUN occupancy (Figure 4J-L). In contrast, SOX2 and c-JUN exhibited no significant binding overlap (Figure 4J-L). Together, these binding analyses demonstrate that SOX21 promotes an anti-tumorigenic gene expression profile by targeting AP-1 bound chromatin regions.

### SOX21 Represses AP-1 Activated Gene Expression in GSCs

Given the central role of AP-1 transcription factors in the maintenance and progression of various types of cancer stem cells, we next investigated the gene regulatory interplay between SOX21 and AP-1. A comparison of genes bound by SOX21 (closest transcriptional start sites) and those regulated by SOX21 revealed that SOX21 predominantly functions to repress targeted genes in GSCs (Figure 5A), which is in contrast to SOX2 that is mainly associated with genes activated by its overexpression (Figure 5A). Interestingly, by comparing gene expression changes induced by pharmacological AP-1 inhibition (via the small molecules T-5224^27^ and SR 11302^28,29^) with those repressed by SOX21, we observed a strong overlap among genes suppressed by AP-1 inhibition and those downregulated by SOX21 induction, which was not found among the genes activated by AP-1 inhibition (Figure 5B, C; Supplementary Figure S7A, B). Consistent with this gene regulatory overlap, gene-set enrichment analysis (GSEA) showed a positive enrichment of “*P53 targets*” and a reduction of “*E2F targets* ” in GSCs treated with either AP-1 inhibitors or DOX-induced SOX21 expression (Figure 5D; Supplementary Figure S7 C, D).

**Figure 5.**
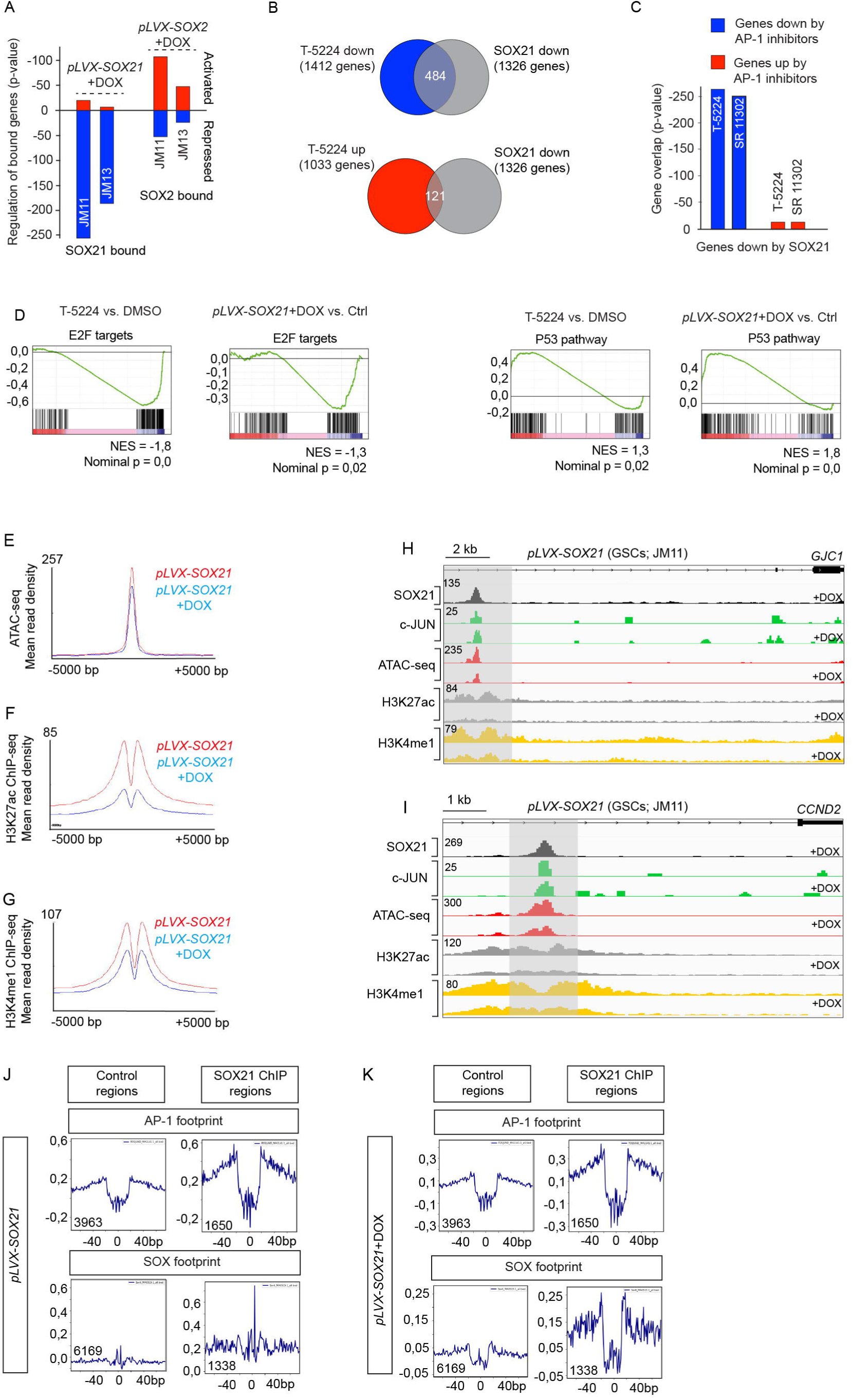
SOX21 Mediated Epigenetic Repression of AP-1 Targeted Genes. (A) Bar graph showing the proportion of transcriptionally activated and repressed SOX21- and SOX2-bound genes after DOX-induced SOX21 or SOX2 expression in JM11 and JM13 GSCs. Up-Quantifications are based on p-values and compared with non-DOX conditions. (B, C) RNA-seq analysis revealing a significant overlap between genes downregulated by SOX21 and AP-1 inhibitors (T-5224 and SR 11302) (represented in blue; B, C), but not among genes upregulated by AP-1 inhibitors (represented in red; B, C) in JM11 GSCs. (D) Gene Set Enrichment Analysis (GSEA) of RNA-seq data comparing the effects of AP-1 inhibition (T-5224) and SOX21 expression on E2F and P53 target genes in JM11 GSCs. (NES, Normalized Enrichment Score. (E-G) Read mean density of ATAC-seq (E), H3K27ac ChIP-seq (F), and H3K4me1 ChIP-seq (G) mapped onto SOX21 and c-JUN bound regions in control (red line) and SOX21 induced (blue line) JM11 GSCs. (H, I) Read density tracks show representative examples of peaks derived from SOX21 ChIP-seq (black), c-JUN ChIP-seq (green), ATAC-seq (red), H3K27ac ChIP-seq (gray), and H3K4me1 ChIP-seq (yellow) in control or SOX21 expressing GSCs. (J, K) Footprint analysis of ATAC-seq data at AP-1 and SOX motifs, present in control or SOX21 bound chromatin regions. Mean aggregate ATAC-seq signals are shown on y-axes and the number of analyzed motifs is stated in the graphs.

The observation that SOX21 represses AP-1-driven gene expression, we next sought to identify the underlying molecular mechanisms. To address this issue, we employed ATAC-seq^30^ to examine how SOX21 induction for 48 hours alters chromatin accessibility at SOX21-and c-JUN-targeted regions. We also performed ChIP-seq experiments to assess their association with the epigenetic markers H3K4me1 and H3K27ac, indicative of primed and active enhancers, respectively^31,32^. Quality control measures confirmed the reliability and reproducibility of the ATAC-seq experiments (Supplementary Figure S8A-E). Principal component analysis of ATAC-seq data showed that DOX-induced SOX21 expression significantly altered the chromatin architecture, separating SOX21-induced samples from controls (Supplementary Figure S8B). We identified approximately 50,000 chromatin regions with altered accessibility following SOX21 induction, with nearly equal numbers exhibiting increased or decreased ATAC-seq signals (Supplementary Figure S8E; Supplementary Table S5). Notably, of the common chromatin regions identified in the SOX21 and c-JUN ChIP-seq data, 95% were accessible in control cells, an accessibility that significantly decreased upon SOX21 induction (Figure 5E, H, I). The reduction in the accessibility of SOX21 and c-JUN targeted chromatin was followed by a strong decrease in H3K27ac and H3K4me1 levels upon SOX21 binding (Figure 5F-I). These findings indicate that the capacity of SOX21 to inhibit AP-1-driven gene expression in GSCs is associated with SOX21-mediated epigenetic inactivation of commonly targeted enhancer regions.

Our findings indicate that c-JUN is present at commonly targeted chromatin regions before SOX21 induction (Fig. 5G, H). To explore how AP-1 binding may influence that of SOX21, we performed transcription factor footprint analysis on ATAC-seq data, a computational method to predict direct transcription factor binding at specific motifs. Comparing SOX21-bound regions with control chromatin regions containing similar SOX and AP-1 motifs, we mostly observed strong enrichment of AP-1 footprints at SOX21-bound enhancers, independent of SOX21 induction (Figure 5J, K). Moreover, despite the widespread presence of SOX motifs in control regions, SOX footprints were mainly found in enhancers that are preoccupied by AP-1 transcription factors. These findings suggest that SOX21 binding and subsequent repression of AP-1-driven gene expression is guided by pre-existing chromatin accessibility and AP-1 occupancy.

### Pharmacological Inhibition of AP-1 Mimics SOX21 Function in GSCs

Given that SOX21 represses AP-1-driven gene transcription, we next explored whether AP-1 inhibition functionally mimics SOX21 activity in GSCs. To address this issue, we examined the effects of AP-1 inhibition on GSC survival, proliferation, and sphere-forming capacity. Inhibition of AP-1 function, with either T-5224 or SR 11302, significantly reduced GSC viability (Figure 6A, B; Supplementary Figure S9A), and suppressed active cell division, as evidenced by decreased EdU incorporation (Figure 6C; Supplementary Figure S9B). Moreover, AP-1 inhibitions compromised GSC self-renewal as indicated by a marked reduction in tumor sphere formation (Figure 6D-F). Thus, inhibition of AP-1 function recapitulates the activity of SOX21 in GSCs. Conversely, overexpression of c-JUN, effectively rescued the SOX21-induced suppression of GSC viability and proliferation (Figure 6G-I; Supplementary Figure S9C, D). Together, these functional experiments confirm that SOX21-mediated suppression of GSC features depends on its ability to repress AP-1 activity.

**Figure 6.**
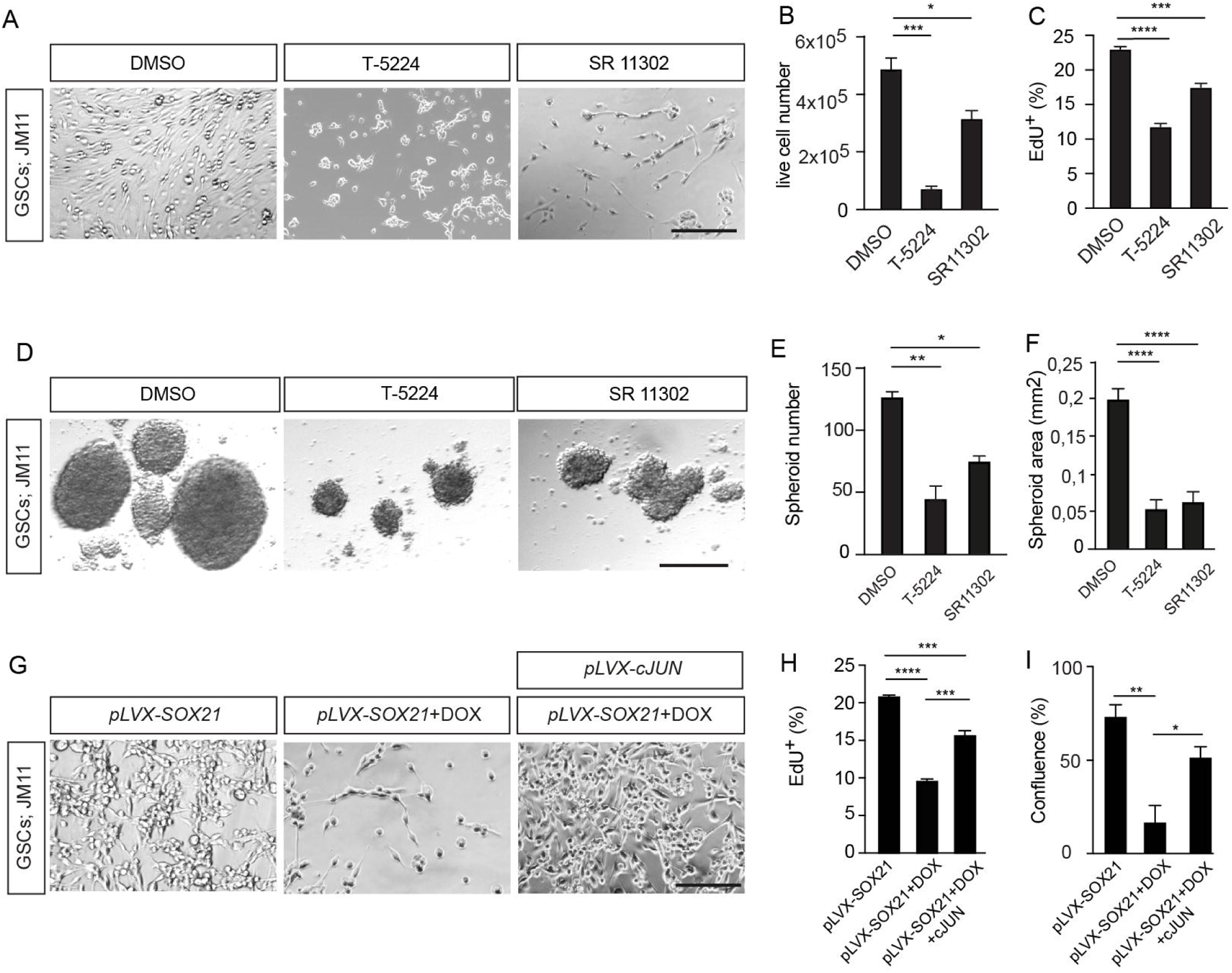
c-JUN Expression Rescues SOX21 Mediated Suppression of GSCs Features. (A) Representative images of GSCs treated with DMSO (control), T-5224, or SR 11302 for 4 days. Scale bar, 50 µm. (B, C) Quantification of cell viability (B) and proliferation (EdU incorporation) (C) following AP-1 inhibition. (D) Sphere formation assay in GSCs treated with AP-1 inhibitors or DMSO. Scale bar, 50 µm. ((E, F) Quantitative assessments of spheroid numbers (E) and spheroid area (F) in response to the treatment with T-5224 or SR 11302. (G) Overexpression of c-JUN for 4 days rescues the reduction in cell number induced by DOX-induced SOX21 expression. Scale bar, 50 µm. (H, I) Quantification of proliferation (EdU incorporation) (H) and viability (plate confluence) (I) following c-JUN overexpression, with or without DOX-induced SOX21 expression.

## Discussion

The prevailing treatment protocol for GBM consists of surgical resection followed by radiation and chemotherapy with DNA-alkylating agents^33^. However, this approach often fails due to its inability to eradicate treatment-resistant GSCs. Therefore, the development of novel therapeutic strategies that simultaneously target both the bulk tumor cells and GSCs is of critical importance.

Previous loss-of-function studies in mice models have demonstrated that SOX21 is essential for preventing malignant transformation of brain neural stem cells exposed to oncogenic insult^15^. Moreover, experiments in which SOX21 is overexpressed in primary human GSCs before transplantation into mice highlight the potential of SOX21 to inhibit their tumor-initiating capacity^15,16^. While these results align with the tumor suppressor function of SOX21 their therapeutic relevance can be questioned. By generating patient-derived GSCs, equipped with an inducible SOX21 expression system, we have in this study demonstrated the ability of SOX21 to impede tumor progression upon its induced expression in established GBM, which ultimately leads to an extended survival of the orthotopically transplanted mice. Our findings reveal a an inverse correlation between SOX21 expression levels and GSC self-renewal and survival. Notably, the reduction in GSC properties following DOX-induced SOX21 expression was accompanied by a significant decrease in the expression of GSC-related genes such as *SOX2*, *HDAC9*, *CDK6,* and *EFNB2*, all of which have been implicated in GBM pathogenesis^21–25^.

A key finding of this study is that SOX21 mediates its tumor suppressor function by repressing the activity of the AP-1 transcriptional factors, which are well-established drivers of cancer stem cell properties, tumor aggressiveness, and treatment resistance^9,34,35^. This conclusion is supported by the identification of a physical interaction between SOX21 and c-JUN in GSCs, as well as the strong overlap between c-JUN and SOX21 targeted chromatin regions. Moreover, pharmacological inhibition of AP-1 in GSCs phenocopied the effects of SOX21 induction, including the downregulation of key genetic pathways and a reduction in GSC proliferation, viability, and tumor-sphere formation. Conversely, forced c-JUN expression rescued SOX21 repressed GSC viability and proliferation. Interestingly, while high SOX21 expression levels correlate with improved GBM patient outcomes, elevated levels of AP-1 transcription factors generally predict poorer prognosis^9^.

How is then SOX21 counteracting the AP-1 function in GSCs? We show that SOX21 binding is preferentially directed to chromatin regions that are accessible and pre-bound by AP-1. AP-1 transcription factors promote chromatin opening through an interaction with the SWI/SNF chromatin remodeling complex^35–37^, a function that can contribute to tumor initiation^35^. In contrast, the binding of SOX21 leads to reduced accessibility of targeted chromatin and a strong decrease in the presence of H3K27ac and, to a lesser extent, of H3K4me1, indicating a transition from an active to an inactive or poised enhancer state^32^. Hence, SOX21 may suppress GSC maintenance and proliferation by limiting the capacity of AP-1 to facilitate enhancer accessibility and thereby decreasing the binding of additional factors, driving tumor-promoting gene expression.

In contrast to SOX21, SOX2 has been referred to as an oncogenic factor due to its essential role in the maintenance and expansion of cancer stem cells ^11,25,38–42^. In gliomas, elevated SOX2 expression correlates with increased tumor aggressiveness and poor patient prognosis^43^. Although induced SOX2 expression did not alter GBM growth pattern in our model, previous knock-down studies have confirmed its necessity for GSC proliferation and tumor-inducing capacity^24,25^. Structurally, the DNA-binding high-mobility group (HMG) domains of SOX21 and SOX2 are highly conserved; however, while SOX21 functions as a transcriptional repressor, SOX2 acts as a transcriptional activator^44^. Interestingly, a modified version of SOX2 version, engineered to function as an epigenetic repressor, exhibits similar tumor-suppressive properties as SOX21, effectively counteracting GBM progression when delivered via viral vectors into the brains of mouse tumor models^45^. Although one interpretation of these findings aligns with the previous idea that SOX21 and SOX2 act on a shared set of target genes^14,46^, we found a surprisingly limited target site overlap (< 10%) in GSCs. Consequently, induced SOX21 expression regulated an insignificant fraction of the SOX2-bound genes in GSCs. One explanation for the divergent binding pattern of SOX21 and SOX2 is that the target selection of SOX proteins is generally dependent on their collaboration with partner transcription factors^46^. Here we demonstrated that while SOX21 physically interacts with c-JUN and binds in the vicinity of AP-1 transcription factors, SOX2 binding is enriched around motifs for POU transcription factors, which are well-established partner factors of SOX2 in different stem cell compartments^46–49^. These findings suggest that although SOX21 and SOX2 exert opposing effects on similar processes in GSCs, their specific binding patterns and cooperation with unique partner factors indicate that they achieve these functions in GSCs by controlling distinct molecular pathways.

This study not only identifies SOX21 as a key tumor suppressor in GSCs through its repression of AP-1-driven oncogenic pathway but also highlights its potential therapeutic relevance. While direct pharmacological activation of SOX21 may be challenging, targeting its associated or downstream pathways represents a promising strategy. By elucidating the molecular mechanisms by which SOX21 modulates glioma biology, our findings provide valuable insights into GSC regulation that could inform the development of innovative therapies aimed at preventing the recurrence of GBM.

## Supplementary material

Supplementary material is available online at Neuro-Oncology (https://academic.oup.com/neuro-oncology).

## Ethics

All animal procedures and experiments were performed in accordance with Swedish animal welfare laws authorized by the Stockholm Animal Ethics Committee: Dnr 3796-2020. All analyses of human material were performed in accordance with an authorization issued by the Swedish Ethical Review Authority: Dnr 2024-06662-02.

## Authorship statement

Conceptualization, J.M., M.B., E.R., and J.Y.; methodology, E.R., J.Y., M.B., and I.K.; formal analysis, E.R.., J.Y., and G.B.; investigation, E.R., J.Y., I.K., V.M., G.B., and M.B.; resources, J.M., O.P.; data curation, E.R., J.Y., V.M., and G.B.; writing original draft, E.R.; J.Y, and J.M.; visualization, E.R., and J.Y.; supervision, J.M., I.K., and M.B.; funding acquisition, J.M.

## Acknowledgement

We are grateful to Dr. Behzad Yaghmaeian for valuable bioinformatic guidance. We would like to thank Dr. Johan Holmberg, Dr. Johan Ericson for helpful discussions and for critically reading the manuscript. This study was made possible in part to The National Genomics Infrastructure in Stockholm (funded by Science for Life Laboratory, the Knut and Alice Wallenberg Foundation, and the Swedish Research Council) and NAISS/Uppsala Multidisciplinary Center for Advanced Computational Science that for assisted us with massively parallel sequencing and enabled access to the UPPMAX computational infrastructure.

