## Supplementary Info for "SOX21 Suppresses Glioblastoma Growth by Repressing AP-1 Activity"

### **Supplementary Materials and Methods**

#### **ChIP-Seq**

ChIP experiments were conducted following established protocols using the following antibodies: goat anti-SOX21, rabbit anti-SOX2, rabbit anti-H3K4me1, and rabbit anti-H3K27Ac. ChIP libraries were prepared using the ThruPLEX DNA-Seq Kit (Takara) and sequenced on the Illumina NovaSeq6000 SP-100 platform, generating paired-end 2x50 bp or 2x150 bp reads, with a sequencing depth exceeding 2 ×10^7 reads per experiment.

ChIP reads were aligned to the human genome assembly hg19 using Bowtie (v1.3.1). Correlation plots between replicate BAM files were generated with deepTools (v2.5.1). MACS2 was used for peak calling under default settings, except for SOX2 and SOX21 ChIP in JM13, where an FDR threshold of 0.001 was applied. Consensus peak sets were defined by detecting signals across all three replicates. Blacklisted regions from hg19 (Encodeproject.org) were excluded. Centrally enriched motifs were identified with CentriMo, and spacing between DNA-binding motifs was analyzed with SpaMo (v5.5.5).

For H3K27Ac and H3K4me1 ChIP-seq, reads were pooled from two replicates each. After duplicate removal, pooled files were used without further processing. To compare ChIP-seq data with ATAC-seq (mapped to hg38), peak regions were converted from hg19 to hg38 using LiftOver (UCSC). Peak overlaps (>50%) were considered significant using the BEDTools intersect function. SeqMINER was used to map H3K27Ac-ChIP, H3K4me1-ChIP, and ATAC-seq reads onto SOX21 peak regions.

Gene annotation was performed using GREAT (v3.0.0). Motif enrichment was analyzed using HOMER (findMotifsGenome.pl, default settings), with peak sets serving as reciprocal background controls in two separate HOMER runs.

#### **ATAC-seq**

ATAC-seq was performed using three replicates per condition, with 50,000 cells per replicate. Tn5-based ATAC experiments were conducted at SciLife Lab (Stockholm, Sweden), with sequencing performed on an Illumina NextSeq2000, generating paired-end 2x50 bp reads. FastQ reads were processed using the nf-core ATAC-seq bioinformatics pipeline (NGI, SciLife Lab, v2.1.2) for mapping, quality control, peak calling, and annotation. Peak differences were analyzed with DiffBind. Footprint analysis on regions less accessible upon SOX21 induction was performed using TOBIAS after Tn5 insertion bias correction. ATACorrect was used for global bias correction, and BINDetect and ScoreBigwig were used to normalize subsets of SOX21 ChIP-seq peaks.

#### **RNA-seq**

Approximately 200,000 cells per sample were harvested 45-48 hours post-DOX treatment or control conditions. RNA was extracted using the RNeasy Prep Kit (Qiagen) following manufacturer guidelines. RNA-seq libraries were generated with the TruSeq RNA Library Prep Kit v2 (Illumina) and sequenced on a NovoSeq6000, producing 2x150 bp reads. FastQ files underwent demultiplexing, quality control, and alignment to hg38 using the NGI SciLife Lab RNA-seq pipeline (v3.14.0). Differential gene expression analysis was conducted with DESeq2, identifying significantly regulated genes with log2 fold change < -1 or >1 and adjusted p-value < 0.05. Volcano plots were generated using the EnhancedVolcano R package. Heatmaps of RNA-seq read counts for selected Gene Ontology (GO) gene sets were created using the R heatmap function. GO term enrichment was visualized with ToppGene.

Isomap analysis, a nonlinear dimensionality reduction technique, was implemented using sklearn.manifold.Isomap, with parameters set to 64 neighbors (preserving local and global structures) and two components (for visualization). Geodesic distances were computed to project high-dimensional data into a two-dimensional space. TPM for JM11, JM12, and JM13 were analyzed to evaluate clustering relative to 165 glioblastoma TCGA samples and 44 glioblastoma cell line data sets^1,2^. Normalization of datasets was performed using two approaches: Z-score normalization (mean of 0, standard deviation of 1) and quantile normalization (NPN)^3^. Datasets for JM11, JM12, JM13, glioblastoma control cells, and TCGA samples were aligned by shared gene names, removing genes with excessive missing values to maintain consistency during dimensionality reduction.

#### **Antibodies**

Primary antibodies: SOX21 (R&D, AF3538, 1:150); SOX2 (Seven Hills Bioreagent, WRAB-1236, 1:500); HuNu (Abcam, ab190710, 1:100); Ki67 (ThermoFisher, 14-5698-82, 1:200; Abcam, ab16667, 1:250); FLAG (Abcam, ab205606, 1:200); HA (Santa Cruz, sc-7392, 1:200); H3K27Ac (Diagenode, C15410174); H3K4me1 (Diagenode, C15410194).

Secondary antibodies: Donkey anti-Goat-488 (Alexa Fluor Invitrogen, A11055); Donkey anti-Rat-488 (Alexa Fluor Invitrogen, A2120); Donkey anti-Rabbit-555 (Alexa Fluor Invitrogen, A31572); Donkey anti-Mouse-555 (Alexa Fluor Invitrogen, A31570); Donkey anti-Goat-555 (Alexa Fluor Invitrogen, A21432); Donkey anti-Rabbit-488 (Alexa Fluor Invitrogen, A21206).

#### **Data Availability**

Raw fastq files from ChIP-seq, RNA-seq, and ATAC-seq experiments (240 files) are available via BioProject ID PRJNA1128363:

[NCBI BioProject PRJNA1128363](http://www.ncbi.nlm.nih.gov/bioproject/1128363).

**Supplementary Figure legends**

**Supplementary Fig. S1** SOX21 Expression is Confined to Self-Renewing Stem Cells.

(A-D) Immunohistochemical analysis of GBM surgical specimens (GBM1705 and GBM1405) reveals that SOX21 protein is predominantly localized in SOX2⁺ stem-like cells and is present in proliferating KI67⁺ cells. (B, D) Bar graphs depict the statistical co-expression analysis of SOX21, SOX2, and KI67 in these GBM samples. Scale bar: 50 µm in (A, C).

**Supplementary Fig. S2** Molecular Subtype Specification of Primary GSCs.

Isometric mapping (Isomap) analysis of RNA-seq data from GSC lines (JM11, JM12, and JM13; replicates #1–3) and 209 GBM tissue samples of known molecular subtypes classifies JM11 as Classical, JM12 as Proneural, and JM13 as Mesenchymal.

**Supplementary Fig. S3** Validation of Inducible SOX21 and SOX2 Expression Systems.

(A) Western blot analysis showing FLAG-tagged SOX21 protein levels in JM12 GSCs transduced with either a control vector (*pLVX*) or an inducible SOX21 expression system (*pLVX-SOX21*), cultured with or without DOX for 48 hours. (B, C) Immunohistochemical staining demonstrating the efficiency of FLAG-tagged SOX21 (B) and SOX2 (C) induction in GSCs (JM11–JM13) following 48 hours of DOX treatment. Scale bar: 30 µm in (B, C).

**Supplementary Fig. S4** Impact of Induced SOX21 and SOX2 Expression on GBM Progression in Mice.

(A) Bioluminescence imaging confirms tumor establishment before DOX treatment in mice transplanted with control or SOX21-inducible JM11 GSCs. (B) Tumor growth curves show luciferase-based quantification of tumor burden in mice receiving either control or DOX-supplemented food. (C, D) Kaplan-Meier survival analysis demonstrates a significant survival advantage in mice with SOX21-inducible JM11 (C) and JM12 (D) GSCs. (E) Representative H&E-stained end-state tumor from a DOX-treated mouse transplanted with control GSCs (JM13). Scale bar: 1.5 mm. (F, G) High-magnification images of immunohistochemical staining for human nuclei (HuNu), FLAG-tagged SOX21, SOX21, and SOX2. Scale bar: 75 µm. (H) Quantification demonstrating a reduction of KI67⁺ proliferative cells in SOX21-expressing tumors. (I) Bioluminescence imaging confirms tumor establishment before DOX treatment in mice transplanted with control or SOX2-inducible GSCs (JM11). (J, K) No significant differences in tumor growth (J) or survival (K) were observed upon DOX-induced SOX2 expression. (L) Bioluminescence imaging confirms tumor establishment before DOX treatment in mice transplanted with control or SOX2-inducible GSCs (JM12). (M, N) No significant differences in tumor growth (M) or survival of transplanted mice (N) could be detected after DOX-induced SOX2 expression.

**Supplementary Fig. S5** Gene Regulation by DOX and SOX2 in GSCs.

(A, B) Volcano plots display differentially expressed genes in control GSCs (JM11 in A, JM13 in B) carrying pLVX (without SOX21), cultured with or without DOX for 48 hours. Upregulated genes (red) and downregulated genes (blue) are indicated (false discovery rate < 0.01). (C) Western blot analysis aligns with transcriptome data and shows a substantial upregulation of P21 protein in JM11 GSCs following DOX-induced SOX21 expression. (D) Heatmaps show the regulation of gene sets and their linked GO-terms in control GSCs transduced with *pLVX*, cultured with or without DOX for 48 hours. (E, F) Volcano plots showing differentially expressed genes in JM11 (E) and JM13 GSCs, following 48 hours of DOX-induced SOX2 expression. Genes up- and downregulated, in comparison to control cells cultured without DOX (*pLVX-SOX2*), are represented with red and blue dots, respectively (false discovery rate < 0,01). (G) Heatmaps show gene sets and their linked GO-terms deregulated upon DOX-induced SOX2 expression. (H) GSEA of differentially regulated genes (blue bars down- and red bars upregulated) in JM11 and JM13 GSCs following SOX2 induction (*pLVX-SOX2*+DOX *vs.* *pLVX-SOX2*).

**Supplementary Fig. S6** Characterization of AP-1 Motifs in SOX21 and SOX2 Targeted Regions.

(A) Heatmap with an associated dendrogram shows the clustering and cell-type specific similarities between SOX21 and SOX2 ChIP-seq replicates in JM11 and JM13 GSCs. (B) Venn diagrams comparing SOX21 ChIP-seq peak regions in JM11 and JM13 GSCs, based on three independent replicates. (C, D) Venn diagrams comparing SOX21 and SOX2 ChIP-seq peak regions in JM11 (C) and JM13 (D) GSCs. (E, F) Centrally enriched SOX motifs in SOX21 (E) and SOX2 (F) ChIP-seq peaks. Solid lines represent data from JM11 and dashed lines data from JM13. P-values of most significant SOX motifs are shown. (G, H) Enrichment of centrally positioned SOX motifs (black) and AP-1 motifs (green) in SOX21 (G) and SOX2 (H) ChIP-seq peaks in JM13 GSCs, with corresponding p-values. (I, J) GSCs overexpressing c-JUN (I), and SOX21 (J) stained with SOX21 and c-JUN antibodies. Results confirm the specificity of the SOX21 antibodies used in the SOX21 ChIP-seq experiments. Scale bar: 30 µm. (K, L) Graphs depicting the most significant motif spacing between SOX and AP-1 motifs in SOX21 ChIP-seq peaks (K) and between SOX and POU motifs in SOX2 ChIP-seq peaks (L). Distance calculations are based on nucleotide spacing between the last base of the SOX motif and the first base of the AP-1 (K) or POU (L) motif.

**Supplementary Fig. S7** Comparative Functions of AP-1 Inhibitors and SOX21 in GSCs.

(A, B) RNA-seq analysis revealing a significant overlap between genes downregulated by SOX21 and AP-1 inhibitors (T-5224 and SR 11302) (represented in blue; B, C), but not among genes upregulated by AP-1 inhibitors (represented in red; B, C) in JM13 GSCs. (C, D) GSEA analysis of RNA-seq data from AP-1 inhibitor T-5224 (C) and SOX21 (D) expression in JM13 GSCs, focusing on E2F targets and P53 targets.

**Supplementary Fig. S8** Quality Control Measures for ATAC-seq Experiments.

(A) Bar graph shows the distribution of genomic features across ATAC-seq peak regions. (B) Principal component analysis of ATAC-seq read data, displays similarities between accessible chromatin regions in control and SOX21-expressing GSCs. Replicates of each experiment are encircled. (C, D) Heatmaps with associated dendrograms show similarities between ATAC-seq peaks in GSCs harboring *pLVX* and *pLVX-SOX21*, cultured with or without DOX for 48 hours. (E) MA-plot showing changes in chromatin accessibility, identified by ATAC-seq, after 48 hours of DOX-induced SOX21 expression (*pLVX-SOX21*+DOX). Regions significantly increased and decreased in their accessibility compared to control cells (*pLVX-SOX21*) are marked with blue dots (false discovery rate < 0.05).

**Supplementary Fig. S9** Overexpression of c-JUN Rescues SOX21 Mediated Growth Suppression in GSCs.

(A) Representative images show JM13 GSCs treated with DMSO (control), T-5224, or SR 11302 for 4 days. Scale bar: 60 µm. (B) Quantitative assessments of proliferation (EdU incorporation) in response to the treatment with T-5224 or SR 11302. (C) Overexpression of c-JUN in GSCs (JM13) counteracted the cell reduction caused by DOX-induced SOX21 expression. Scale bar: 60 µm. (D) Bar graph shows a quantitative assessment of proliferation (EdU incorporation) in response to DOX-induced SOX21 and c-JUN expression, demonstrating that high levels of c-JUN counterbalance the reduced proliferation caused by SOX21.
